# Cryo-EM structure of the FtsH periplasmic domain reveals functional dynamics

**DOI:** 10.1101/2025.10.03.679995

**Authors:** Günce Göc, Sathish K. N. Yadav, George Orriss, Ufuk Borucu, Imre Berger, Christiane Schaffitzel, Burak V. Kabasakal

## Abstract

FtsH, an essential AAA+ metalloprotease, maintains cellular homeostasis by degrading misfolded and membrane-associated proteins. Here, we report cryo-EM structures of the *Escherichia coli* FtsH periplasmic domain (FtsH-PD) revealing insights into its conformational flexibility. Initial 2D class averages suggested three distinct orientations, right-handed and left-handed maps of FtsH-PD, and a map with a different conformation. The 4.9 Å structure of FtsH-PD exhibits the conserved α+β fold, while the 7.3 Å map with the different conformation displays a 20º clockwise rotation of two alpha helices. These findings support a model where conformational changes are present not only in the FtsH cytosolic domain, but also in the periplasmic domain and potentially facilitate substrate translocation through a combination of mechanisms involving both the FtsH-PD and the HflKC complexed with FtsH, along with lipid-scramblase activity to assist in membrane protein extraction. This study points out novel perspectives on how conformational changes in the periplasmic domain contribute to FtsH substrate degradation mechanisms.

## Introduction

Proteases with diverse cellular activities associated with ATP *(AAA+; ATPases associated with diverse cellular activities)* are conserved in all organisms and function as large hexameric or heptameric oligomers (Vostrukhina et al., 2015). AAA+ Proteases play a key role by being involved in a multitude of metabolic activities, including degradation of misfolded or unfolded proteins, DNA replication, membrane fusion, and signal transduction. The degradation of protein substrates by AAA+ proteases occurs in two stages, regarding the two domains of AAA+ Proteases: the ATPase domain provides the mechanical energy derived from the conversion of chemical energy using ATP, which is then employed to unfold the three-dimensional structure of the target protein in the proteolytic domain. Subsequently, the degraded target protein is decomposed into polypeptide chains and amino acids (Yedidi et al., 2017).

Degradation process of the target proteins represents a crucial mechanism for the maintenance of protein homeostasis, which is why it is referred to as protein quality control. As the most significant member of the protein quality control mechanism with considerable substrate variability, FtsH *(Filamentation Temperature Sensitive Protein H)* plays a pivotal role in maintaining cellular homeostasis as a AAA+ metalloprotease. The uniqueness of FtsH comes from being an integral membrane protein itself and its ability to degrade both membrane-bound and soluble proteins. Consequently, FtsH is capable of degrading a wide range of substrates, thereby facilitating the involvement of various metabolic processes (Ito & Akiyama, 2005; Kato & Sakamoto, 2018; Yi et al., 2022).

As a universally conserved hexameric AAA+ metalloprotease, each *Escherichia coli* FtsH subunit has a cytosolic domain (FtsH-CD) with ATPase and proteolytic activity and a periplasmic domain (FtsH-PD) in a single polypeptide chain (Figure 1) (Bittner et al., 2017; Ito & Akiyama, 2005). The N-terminal part comprising two transmembrane helices is followed by the AAA+ ATPase domain which contains Walker A (amino acid sequence: GPPGTGKT) and Walker B motifs (amino acid sequence: VAGCDE). These motifs coordinate binding and hydrolysis of ATP, with the participation of bound Zn^2+^ and H_2_O. To provide mechanical energy for substrate translocation hydrolysis of ATP is necessary. ATP hydrolysis results in a “pulling” which facilitates conformational changes in the cytoplasmic domain of FtsH, causing the substrate to move forward through the central cavity of the proteolytic domain, leading to degradation of the bound substrate into small oligopeptides (Carvalho et al., 2021; Sauer & Baker, 2011).

**Figure 1.**
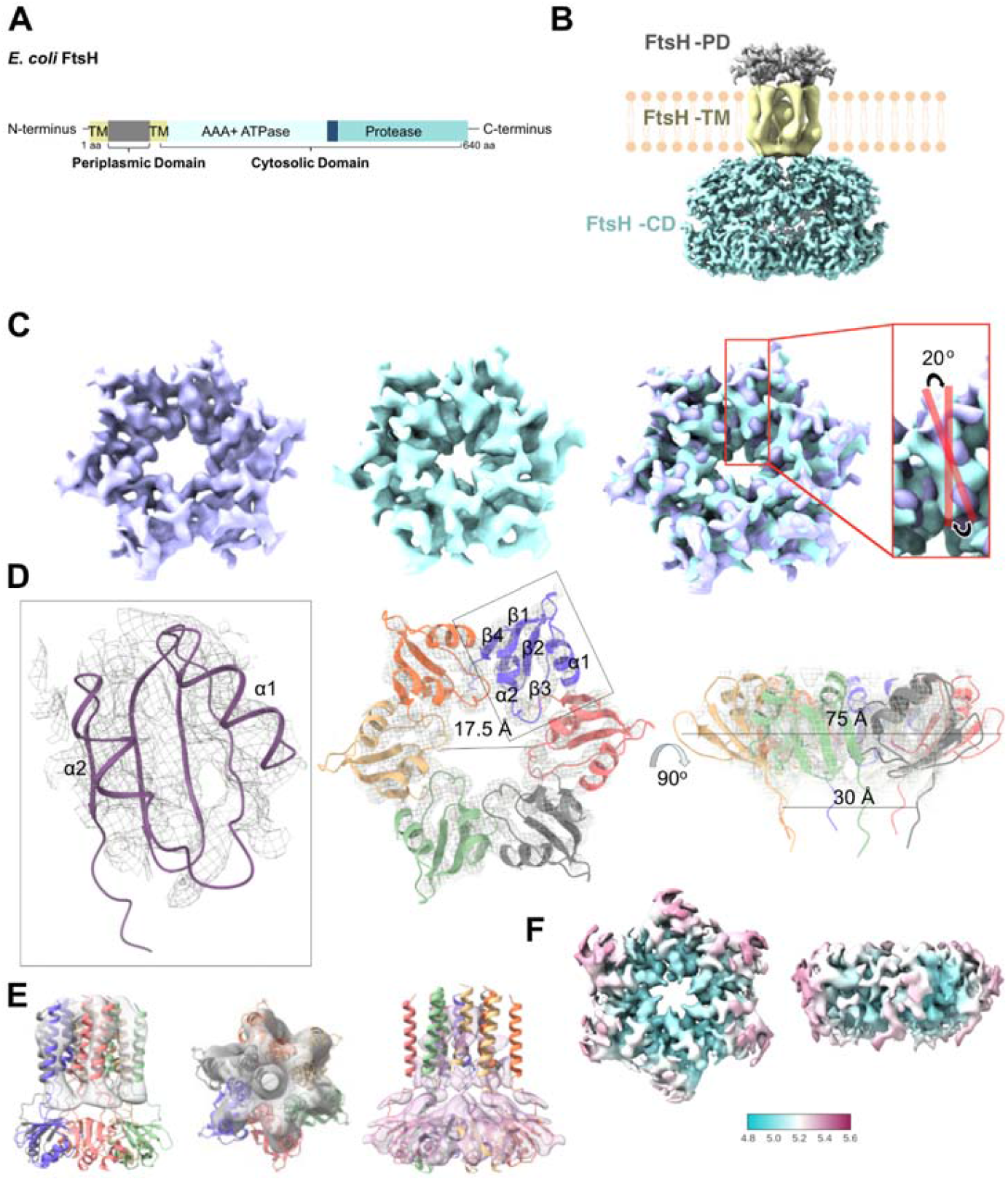
Structural Analysis of *E. coli* FtsH. **A. Domain architecture and structure of *E. coli* FtsH domains**. Schematic representation of the domain architecture of *E. coli* FtsH. The protein comprises an N-terminal periplasmic domain (grey), two transmembrane helices (yellow), a cytosolic AAA+ ATPase domain (light blue), and a protease domain (turquoise). **B. Structural model of the FtsH complex integrated into the membrane**. The periplasmic domain (FtsH-PD, grey), transmembrane helices (FtsH-TM, yellow), and cytosolic domains (FtsH-CD, turquoise) are depicted, based on the maps from this study (PD and TM) and the EMD-32521 (CD) (Qiao et al., 2022). **C. Counterclockwise (CCW) (purple) and novel orientation (NO) (turquoise) maps of FtsH-PD**. Superposition of maps shows that FtsH-PD undergoes a conformational change by a 20º clockwise rotation, highlighted by sticks on the α2 helices. **D. Hexameric model of the FtsH-PD**. Atomic model fitted into the cryo-EM map (grey mesh). One subunit is highlighted (box). Alpha helices undergoing a conformational change are labelled as α1 and α2. **E. FtsH-TM maps**. AlphaFold2 model of the hexameric FtsH-PD-TM fitted into FtsH-TM maps reconstructed from subtracted particles (grey), with a loose mask (salmon). **F. Local resolution map of FtsH-PD**, indicating a resolution ranging from 4.8 to 5.6 Å.

Structures of ADP-bound FtsH from *Thermotoga maritima* (Liu et al., 2022) and *Aquifex aeolicus* (Carvalho et al., 2021), and an ATP-bound yeast homolog YME1 structure have been determined (Puchades et al., 2017). The crystal structures of the FtsH-CD indicated that the ATPase domain can exhibit C3 (Suno et al., 2006), C2 (Bieniossek et al., 2006), and C6 symmetry (Bieniossek et al., 2009b), adopting? different conformations depending on the identity of the bound nucleotide (ATP, ADP), or lack thereof (apo structure). This nature of these conformational changes has led to the current consensus that the “pulling” mechanism is most probable (Bieniossek et al., 2009b). In addition to crystal structures, a cryo-EM structure of the FtsH-CD homologue, YME1 (pdb 6az0), exhibited a “staircase” symmetry of the ATPase domain (Puchades et al., 2017). This structure indicates an opening of 1.4 nm, which would be sufficient for the passage of small peptides. Comparison of mitochondrial inner membrane protease YME1-CD with the apo-state structure of *T. maritima* FtsH-CD (pdb 2ce7) shows that the ATPase domain is capable of rotating 28° to facilitate the access of peptides to the central pore, further supporting the “pulling” hypothesis (Puchades et al., 2017; Carvalho et al., 2021). There is no high resolution full-size FtsH structure reported to date. Therefore, the effect of FtsH cytosolic domain conformational changes on the periplasmic domain of FtsH, or possible conformational changes in the periplasmic domain remain unclear.

Here, we report the structural characterization of *E. coli* FtsH-PD, that we obtained from the purification of FtsH-HflKC complexes using electron cryo-microscopy (cryo-EM). The FtsH-PD exhibits C6 symmetry. Our cryo-EM analysis revealed two different orientations of FtsH-PD, named counterclockwise (CCW, EMD-66269) and a novel orientation (NO, EMD-66013). These conformations were resolved to resolutions of 4.9 Å and 7.3 Å, respectively. The NO map exhibited a 20º clockwise rotation of two helices relative to the FtsH-PD-CCW map. The two conformations suggest a molecular mechanism how the periplasmic domain of FtsH may be involved in the proteolysis of membrane-bound substrates.

## Materials and Methods

### Protein Expression and Purification Conditions

Protein expression was carried out by using one single plasmid carrying genes encoding FtsH-3×StrepTag and HflKC-10×His, each under the control of inducible promoters; arabinose, and T7, respectively, which was generated using the ACEMBL system (Bieniossek et al., 2009a). *E. coli* C43 (DE3) cells expressing the C-terminal streptactin-tagged FtsH-HflK-HflC were grown overnight at 37°C, shaking with 100 μg/ml Ampicillin, 50 μg/ml Spectinomycin in Luria-Bertani broth (LB) media. The expression media (2xYT) was inoculated at 1:50 dilution, incubated at 37°C, shaking until OD at 600nm reaches 0.6-0.8. Protein expression was inducted with 1mM Isopropyl-β-D-thiogalactopyranosid (IPTG), and 0.2% (w/v) arabinose at 37°C for 3 hrs. 0.2 mM ATP was added to the media at the time of induction. At the end of three hours, cells were harvested by centrifugation at 4500 xg for 25 min. The separated cells were resuspended in Buffer A (20 mM HEPES, pH 8.0, 150 mM KCl, 5 mM beta-mercaptoethanol, 20% (v/v) glycerol). Protease inhibitor tablets were added to resuspended cells, and cells were lysed using a cell disrupter at 25 kpsi by passing twice. Five units/ml of benzonase and RNAse were added to lysed cells, and spinned at 20 000 xg for 40 min at 4°C to discard pellets. Membranes were isolated by centrifuging the supernatant at 100 000 xg using Type 45 Ti rotor (Beckman Coulter) for 160 min at 4°C. Membranes were resuspended in Buffer A using a glass Dounce homogeniser, and solubilized in 2% (w/v) n-dodecyl-β-D-maltoside (DDM) (Sigma) by gently shaking for 2 hrs at 4°C. Supernatant containing the membrane proteins were obtained by centrifugation at 100 000 xg using Type 45 Ti rotor for 160 min at 4°C, and loaded onto streptactin resin equilibrated with Buffer A’ (20 mM HEPES, pH 8.0, 150 mM KCl, 1 mM beta-mercaptoethanol, 0.01% (w/v) DDM. Proteins were eluted with 4 mM desthiobiotin and concentrated to ∼200 μl using 100 kDa MWCO concentrators.

### ATPase Activity Assay of FtsH & FtsH-HflKC

ATPase activity of membrane protein FtsH was analyzed using EnzCheck™ Phosphate Assay Kit (Thermo Fisher Scientific, Cat. No. E6646). Measurements were conducted according to the manufacturer’s instructions. A standard calibration curve for P_i_ quantification was generated using a series of reaction mixtures with increasing concentrations of potassium dihydrogen phosphate (KH□PO□) as the phosphate standard, ranging from 0 to 100 µM final concentration. All reactions were mixed thoroughly and incubated at 22 °C for 30 minutes. Absorbance was then measured at 360 nm using a UV-3600i Plus UV-Vis-NIR spectrophotometer (Shimadzu, JP). The background absorbance from the phosphate-free control was subtracted from all measurements. A standard curve was generated by plotting the known phosphate concentrations (X-axis) against the corresponding absorbance values (Y-axis), and a linear regression analysis was performed to determine the equation of the trendline, which was subsequently used to quantify Pi generated in experimental samples.

In all ATPase assay conditions, the FtsH and FtsH-HflKC was used at a final concentration of 3□µM, along with a substrate solution containing 5□mM ATP were added to reactions. Following substrate addition, absorbance at 360 nm was measured at time zero and subsequently at 15-minute intervals for a total duration of 60 minutes. Time-dependent increases in absorbance were recorded and converted to phosphate concentrations using the previously generated standard curve, allowing for quantification of ATPase activity.

### Protease Activity Assay of FtsH & FtsH-HflKC

The protease activity of FtsH and FtsH-HflKC was measured using resofurin-labeled 0.4% casein (Sigma-Aldrich, Cat. No. 11 734 334 001) as a substrate, and the measurements were conducted in alignment with the manufacturer’s instructions. Reaction was initiated using 0.5 µM FtsH or FtsH-HflKC and aliquots were taken at definite time points of 15, 30, 45, 60 and 120 minutes. Then reactions were terminated using 5% TCA, followed by 10 minute incubation at 37 °C. Samples were then centrifuged at 12,000 × g for 5 minutes. 0.5 M Tris-HCl (pH 8.8) was added prior to measuring the absorbance at 574 nm.

### Reconstitution in Amphipol

300 μg of protein at 1 mg/ml were solubilized in DDM, and were buffer exchanged into Amphipol (A8-35, Anatrace) at a protein/amphipol mass ratio of 1:7. The protein/amphipol mixture was incubated on a rotator for 2h at 4°C. To remove the excess free detergents, 40X mass excess of SM2 BioBeads (BioRad), previously washed with methanol and water, and dried, added to the protein/amphipol mixture and incubated overnight at 4°C on a rotator. The protein/amphipol/BioBead mixture was centrifuged in a microcentrifuge at 15 000 xg for 15 min. The supernatant containing the detergent-free and amphipol-reconstituted membrane protein was carefully taken, and concentrated in a 0.5 ml concentrator with a 30 kDa MWCO in Buffer C (20mM HEPES, pH 8.0, 150mM KCl) to a final volume of 50 μl. The concentrated protein was then injected to 3.2/300GL Superose 6 column running at 0.05 ml/min of Buffer C, with 0.05 ml fractions collected in a 96-well plate on a micro AKTA.

### Negative stain EM and Cryo-EM sample preparation

Five µL of 0.04 mg/ml protein was applied onto a freshly glow discharged (1 min at 30 mA) carbon film-coated grid (ECF300-Cu Electron Microscopy Sciences), incubated for 1 min, and manually blotted. 5 µl of 3% uranyl acetate was applied onto the same grid and incubated for 1 min before the solution was blotted off. The grid was loaded into a FEI Tecnai12 120 kV BioTwin Spirit TEM. Images were acquired at a nominal magnification of 49,000x with a FEI Ceta camera.

For the Cryo-EM sample, 3 µl of 0.35 mg/ml protein was applied onto a freshly glow discharged (1 min at 30 mA) Quantifoil R1.2/1.3 carbon grid (Agar Scientific), blotted using a Vitrobot MarkIV (Thermo Fisher Scientific) at 80% humidity and 16°C for 1.6 s, and plunge frozen.

### Data Collection and Processing of FtsH-PD and FtsH-TM

A data set with 6331 movies was collected with EPU on a Talos Arctica at GW4 Facility, Bristol, UK with an acceleration voltage of 200 kV and magnification of 130,000x, with an effective pixel size of 0.525 Å (binned by 2) in super-resolution mode on a Gatan K2 Summit direct electron detector and Gatan Quantum GIF energy filter operated in zero-loss mode with a slit width of 20 eV. Movies were collected as 48 frames with a defocus range from -1 to -2 µm and a total exposure of 61.3 e-/Å^2^. Cryo-EM data was processed using the Relion 3.1, 4.0, and 5.0 software packages (Fernandez-Leiro & Scheres, 2017; Zivanov et al., 2020). First, the motion correction of micrographs was performed with MotionCorr2 (Li et al., 2013), and contrast transfer function (CTF) estimation was done using ctffind4.1 (Mindell & Grigorieff, 2003). Micrographs with resolutions better than 4.2 Å were selected for particle selection (5469 out of 6331 micrographs). After two rounds of 2D classification, 929,166 particles were used for subsequent periplasmic and transmembrane region reconstructions. First, particles corresponding to either periplasmic or transmembrane regions, 429,744 and 455,307 particles, respectively, were subtracted (Supplementary Figure S1).

After 3D classification of the periplasmic region subtracted particles, and in parallel 3D-focussed classification rounds using the original extracted particles, three maps with different orientations of periplasmic FtsH were refined; counter-clockwise (CCW) orientation with 9590 particles, clockwise (CW) orientation with 20,776 particles, and novel orientation (NO) with 8990 particles. After ctf and aberration refinement, Bayesian polishing, and post-processing, 5.4 Å map of CCW, 5.2 Å map of CW orientations, and 7.3 Å map of NO FtsH-PD were acquired. As the CW map consisted of left-handed helices, particles of CCW and CW orientations were combined, and re-refined. The final post-processed right-handed map (CCW) reached up to 4.9 Å based on the Fourier Shell Correlation (FSC) 0.143 cutoff (Rosenthal & Henderson, 2003). Local resolution of the final map was determined in Relion 5.0, and visualized with ChimeraX (Supplementary Figure S3) (Meng et al., 2023). DeepEMhancer was used to generate sharpened maps for visualizations and model building (Sanchez-Garcia et al., 2021).

For the transmembrane (TM) region, after 3D classification without applying symmetry with subtracted particles, 349,714 particles were selected. After rounds of 3D classifications (C1 and C6 symmetry), a class with helical features was selected. The final 9.3 Å map of TM region was reconstructed with ctf- and aberration-refined 6499 particles, followed by post-processing (Supplementary Figure S5). Subsequently, the TM region was resolved to 8.2 Å by applying a loose mask on the NO FtsH-PD map (Supplementary Figure S6).

### Structure Modelling of FtsH-PD and FtsH-TM

The FtsH-PD (PDB: 9wus) structure was modelled by molecular replacement using the MolRep in the spherically averaged phased translation function mode within the CCP-EM suite (Burnley et al., 2017). The structural model of the periplasmic domain of FtsH in the FtsH-HflKC structure (PDB: 7wi3) was used as a template (Qiao et al., 2022). One FtsH-PD subunit in the 7wi3 structure was used as an input, and six copies were modelled by molecular replacement. Then, it was refined by the real space refinement tool in Phenix (Liebschener et al., 2019), and Refmac in the CPPEM suite (Murshudov et al., 2011). Manual buildings and adjustments were carried out in Coot (Emsley et al., 2010). The final FtsH-PD model was validated using Phenix validation tools (Supplementary Table S2). As the maps of FtsH-TM region were not sufficient for structural model building, the hexameric predicted model of FtsH, generated by AlphaFold2 (Jumper et al., 2021), run through ColabFold (Mirdita et al., 2022), was fitted into FtsH-TM maps using ChimeraX (Pettersen et al., 2021). Periplasmic and transmembrane regions of the predicted model were used only for fitting.

## Results

### Periplasmic domain of FtsH as a degradation product of FtsH-HflKC overexpression

The periplasmic domain of FtsH (FtsH-PD) was purified and characterised after the overexpression of FtsH-HflKC in *E. coli*: Cell membranes were first solubilized in DDM, and FtsH-HflKC was purified by pulling down via the streptactin-tag on FtsH. The purified complex was then transferred to Amphipol A8-35 and further purified using size exclusion chromatography (Supplementary Figure S2A). SDS-PAGE analyses of the affinity-purified, and Amphipol reconstituted complex both suggested that FtsH undergoes a degradation during the purification (Supplementary Figure S2F). A strong band corresponding to the cytosolic domain of FtsH (∼60 kDa) was persistent during all stages (Supplementary Figure S2F). Additionally, the band corresponding to HflC disappeared during the SE step (Supplementary Figure 2F, shown with an arrow). Consequently, the fraction used for cryo-EM data collection was enriched with the Amphipol-reconstituted periplasmic domain of FtsH, due to FtsH degradation, loss of the FtsH-CD and FtsH-HflKC dissociation. The negative-stain EM grid of this fraction showed uniform circular particles which are ∼10 nm in diameter (Supplementary Figure S2D-E).

### FtsH periplasmic domain reveals two conformations

Cryo-EM analysis of the sample resulted with hexameric star-shape 2D classes with a large Amphipol belt. 2D class averages of FtsH-PD initially suggested three different orientations and conformations. The first is the one with alpha helices oriented at the clockwise (CW) rotation, the second is the one with alpha helices oriented at the counterclockwise (CCW) rotation, and the third is the one with novel orientation (NO) (Supplementary Figure S1). After several particle subtraction, global and focused 3D classification steps, we finally obtained three different maps representing three different orientations. Upon careful consideration, we realized that the map with the CW rotation is the mirror image of the CCW rotated map. Therefore, the CCW map comprised right-handed helices whereas the CW revealed left-handed helices, which is biologically impossible, however possible computationally. The CW map flipped along the z-axis aligned well with the CCW map.

The 4.9 Å CCW FtsH-PD cryo-EM structure (FtsH-PD-CCW) showed the same α + β fold as the NMR (pdb 2muy) (Scharfenberg et al., 2015) and X-ray crystal structure (pdb 4v0b) (Scharfenberg et al., 2015), as well as the periplasmic domain of FtsH obtained within the FtsH-HflKC cryo-EM structures (pdb 7vhp, 7wi3) (Ma et al., 2022; Qiao et al., 2022), comprising two α-helices and five β-strands (Figure 1D, Supplementary Figure S7). In the hexameric structure, alpha helix 1 (α1) and alpha helix 2 (α2) have orientations that exhibit the counterclockwise rotation (Figure1C, purple map).

The third type of map had neither a CW nor a CCW orientation, which we named ‘novel orientation’ (NO). The resolution of this map was 7.3 Å. We could not generate a reliable structural model for this EM map (Figure 1C, turquoise map). It is clear, however, that the α1 and α2 helices undergo a conformational change, adopting a novel conformation. Within the FtsH-PD the α1 and α2 helices undergo a 20□ clockwise rotation compared to the FtsH-PD-CCW structure (Figure 1C, superposed maps, Supplementary Video).

### FtsH transmembrane domain

The cryo-EM map of the FtsH transmembrane domain (FtsH-TM) was reconstructed by using the particles subtracted from the 2D classes corresponding to the transmembrane region. The 9.3 Å map of the computationally isolated FtsH-TM (EMD-66013) revealed hexameric alpha helices. The TM domain is 48 Å in length, 45 Å in diameter (Figure 1E, grey map). The FtsH-TM was also resolved by using a loose mask on the FtsH-NO map, which reached up to 8.2 Å (Figure 1E, salmon map). Two alpha helices in the TM domain were seen in this map. The hexameric AlphaFold2 model of the FtsH-TM and FtsH-PD was fitted in both maps.

### ATP binding may cause conformational changes in the periplasmic FtsH

FtsH is known to adopt different conformations upon ATP binding. Structures of ADP-bound FtsH from *T. maritima* (pdb 7tdo) (Liu et al., 2022), and *Aquifex aeolicus* (pdb 4vv0) (Vostrukhina et al., 2015), and ATP-bound yeast homolog YME1 structure (pdb 6az0) (Puchades et al., 2017) have confirmed this. However, this information is limited to the cytosolic domain of FtsH, where the ATP binding site is located. As there is no high resolution full FtsH structure reported to date, the effect of FtsH cytosolic domain conformational changes on the periplasmic domain remains unclear.

Our hypothesis that ATP binding may cause conformational changes in the periplasmic FtsH is supported by the possibility of the presence of the ADP-bound FtsH in the purified sample. ATP (0.2 mM) was added to the growth medium at the time of induction as we obtained higher cell yields with ATP-enriched media. The cytosolic domain of FtsH, where the ATP binding site is, was not detected in the cryo-EM sample, so ATP binding could not be confirmed structurally.

### Possible scenarios on FtsH degradation mechanism

Hypotheses about how FtsH degrades substrate proteins have been reported in the literature (Bieniossek et al., 2006; Carvalho et al., 2021; Chiba et al., 2002; Murata & Wolf, 2018; Suno et al., 2006; Yamada-Inagawa et al., 2003; Qiao et al., 2022; Ghanbarpour et al., 2025) (Figure 2). The best known hypothesis is that both soluble and membrane protein substrates enter the degradation chamber formed by hexamers through a pore between the cytosolic and transmembrane regions. Cytosolic protein substrates may be recognized by the helical subdomain of the ATPase domains whereas membrane-bound substrates may interact with the FtsH transmembrane region through their transmembrane parts. Substrate recognition is followed by the translocation of both types of substrates into the protease domain for endo-proteolysis. Recent studies proposed new, supporting hypotheses: According to the study by Qiao et al., HflKC may provide an alternative entry route for periplasmic substrates via the pore region located on the top of the FtsH-HflKC complex on the periplasmic side (Qiao et al., 2022). Ghanbarpour et al. suggest that FtsH in the FtsH-HflKC complex has lipid-scramblase activity which is also known as lipid flip-flop. This activity is to facilitate the movement or rearrangement of the membrane lipids across the bilayer. Therefore, the membrane integrity is maintained through this process, and it also facilitates cell growth. It has been reported that this activity also enhances the capacity of FtsH to extract and degrade membrane-bound substrates (Ghanbarpour et al., 2025). Our cryo-EM maps, revealing conformational changes in the periplasmic FtsH structure, agree with these two recent hypotheses. The FtsH periplasmic domain may undergo conformational changes to aid the translocation of periplasmic substrates through the central pore of the periplasmic region of FtsH to reach FtsH’s cytoplasmic ATPase domain. The curved lipid domains in the FtsH-HflKC structures may be indeed related to the degradation of membrane-embedded substrates; where substrate-binding and translocation may induce conformational changes in the transmembrane and periplasmic domains of FtsH.

**Figure 2.**
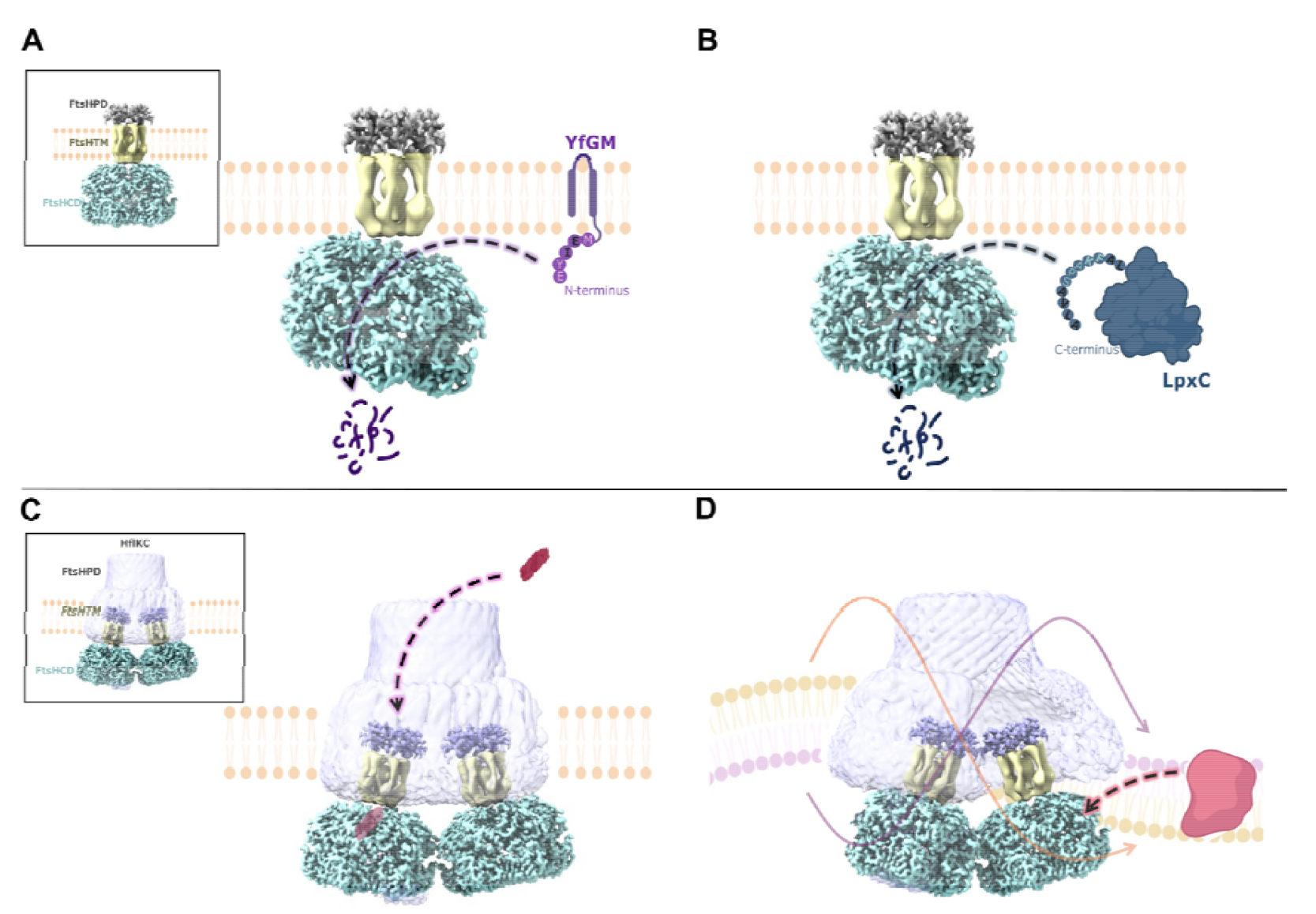
Proposed mechanisms for FtsH-mediated substrate protein degradation. **A. Cytoplasmic interaction of membrane substrates with FtsH**. Recognition and unfolding of membrane substrates, such as YfgM, occur in the cytoplasm. **B. Cytoplasmic interaction of cytosolic substrates with FtsH**. Cytosolic substrates, such as LpxC, are similarly recognized and unfolded in the cytoplasm prior to degradation. **C. Interaction of periplasmic substrates with FtsH**. Periplasmic substrates (indicated in red) are proposed to access FtsH through a gap in the HflKC complex within the periplasmic space, followed by degradation. **D. Lipid scrambling activity by FtsH-HflKC**. Lipid scrambling activity of FtsH-HflKC (purple and orange lipids and arrows) is suggested to facilitate substrate degradation of transmembrane substrates (red). FtsH PD and TM: this study, FtsH CD: EMD-32521 (Qiao et al., 2022), HflKC: EMD-46057 (Ghanbarpour et al., 2025).

## Discussion

FtsH is a universally conserved protease with a large spectrum of substrates, involved in many cellular processes, in particular quality control mechanisms ensuring cell viability. As a membrane-bound protease, FtsH can degrade both cytosolic and membrane proteins. Although FtsH has been studied for decades, structural information is still scarce. Crystal structures of bacterial FtsH (Kryzwda et al., 2002; Suno et al., 2006; Bieniossek et al., 2006; Bieniossek et al., 2009) and recent cryo-EM structures (Ma et al., 2022; Qiao et al., 2022) were reported, but a high-resolution structure of full-length FtsH is still lacking. While crystallization of the whole FtsH is a known challenge, obtaining a good cryo-EM sample of either FtsH or FtsH-HflKC complex is equally challenging (Carvalho et al., 2021; Ma et al., 2022; Qiao et al., 2022). The dynamic nature of FtsH and its auto-cleavage activity need to be overcome to achieve a full-size, atomic resolution FtsH structure. The structure reported here is the first cryo-EM structure of the periplasmic domain of FtsH. To the best of our knowledge, it is the smallest membrane protein reconstruction reported to date using a 200 kV cryo microscope, along with the 3.68 Å structure of multidrug efflux pump in *Mycobacterium tuberculosis* (49 kDa) (Wang et al., 2024) and 3.7 Å structure of glutamate transporter homologue (44 kDa) (Chen et al., 2021).

FtsH research focused on the cytosolic regions in which the proteolytic and ATPase domains of the protein are located. The periplasmic region of FtsH may be involved in substrate recognition, translocation and degradation of proteins in the periplasm. FtsH is capable of degrading membrane-embedded substrates in either the N-to-C or C-to-N direction (Chiba et al., 2000, 2002). It was shown, however, that the proteolysis of the periplasmic domain of the model substrate PhoA by FtsH was halted after degradation of the C-terminal cytosolic region (Chiba et al., 2002). This raises the possibility that the degradation of periplasmic substrates is actually initiated and mediated by the FtsH-HflKC complex through the gap formed by HflKC periplasmic domain. Substrate binding is suggested to be followed by translocation of substrates through the periplasmic region of FtsH, reaching the cytosolic proteolytic domain (Kihara & Akiyama, 1998). Although HflKC has long been accepted as a regulator of FtsH proteolytic activity (Akkulak et al., 2024; Bruckner et al., 2003), FtsH-HflKC was shown to also degrade proteins (Ma et al., 2022; Qiao et al., 2022), aligning with our experimental results (Supplementary Figure S7). A recent study has proposed that the membrane curvature in the FtsH-HflKC complex is related to lipid-scramblase activity. Lipid scrambling would promote the degradation of membrane-embedded proteins by furthering interactions between membrane-bound substrates and the transmembrane region of FtsH (Ghanbarpour et al., 2025). Membrane protein substrates could thus reach the proteolytic active site of the FtsH via the gap formed by the hexameric periplasmic and transmembrane regions of FtsH, furthered by conformational changes in these regions. Conformational change in the FtsH periplasmic region from CCW to NO (Figure 1.C) may be related to membrane-bound substrate movement in the membrane and their degradation. Structural changes, along with the reported scramblase activity in FtsH-HflKC complexes, may aid in substrate extraction by reducing the energy barrier for substrate dislocation. This mechanism is consistent with research suggesting that the degradation of periplasmic substrates such as YfgM requires cytosolic N-terminal recognition (Bittner et al., 2016), implying a coordinated ‘pull and translocate’ process in which initial substrate interaction in the cytosol is followed by unfolding. Notably, the inability of FtsH to degrade the periplasmic domain of PhoA, unless initiated from the cytosolic end, (Chiba et al., 2002) supports a molecular mechanism in which HflKC forms a protected periplasmic substrate channel in its vault-like architecture (Figure 2C).

The retention of proteolytic activity in the FtsH-HflKC complex, as demonstrated in this study and by Qiao et al., (2022) contradicts previous reports suggesting that HflKC is a down-regulator of FtsH proteolytic activity (Kihara et al., 1996; Ito & Akiyama, 2005). Thus, HflKC may act as a topological gatekeeper, limiting substrate access to FtsH, preventing indiscriminate degradation while allowing processive threading of membrane-embedded substrates. This regulatory mechanism is consistent with our observation of dynamic conformational changes in the periplasmic domain (Figure 1C) that may be cooperating with HflKC in directing substrates towards FtsH cytoplasmic active site.

Recent research demonstrating lipid scramblase activity in FtsH-HflKC (Ghanbarpour et al., 2025) suggests a mechanism linking substrate dislocation to membrane remodelling, with asymmetric lipid distribution aiding the extraction of transmembrane domain (TM)-anchored substrates.

The dual role of ATP as a proteolytic activator and destabilizing factor adds complexity for such mechanistic models: For instance, while ATPγS stabilization was critical for structure determination, native ATP hydrolysis likely drives rapid conformational changes in FtsH (>6– 8 ATP molecules per proteolytic event) (Suno et al., 2006). Furthermore, the auto-cleavage ability of FtsH highlights the importance of HflKC in maintaining FtsH and FtsH-HflKC complex integrity during substrate processing, a regulatory mechanism observed in other AAA+ proteases (Bittner et al., 2016).

There are unanswered questions regarding the mechanisms of degradation of membrane-embedded or periplasmic substrates by FtsH alone, or in complex with HflKC. How the periplasmic region of the FtsH is involved in this process requires further research. High resolution structures of full-length FtsH, as well as substrate-bound FtsH and FtsH-HflKC complexes, along with biochemical and functional experiments will enhance our understanding of the molecular mechanisms of FtsH substrate recognition, translocation and degradation. Although further high-resolution studies will be needed to clarify the exact molecular mechanisms underlying this new conformation, this intermediate conformation suggests a dynamic conformational landscape that may be functionally relevant and enriches the known structural repertoire of FtsH.

## Data availability

All cryo-EM maps generated have been deposited to Electron Microscopy Data Bank (EMDB) as entries EMD-66269 (FtsH periplasmic domain), EMD-66013 (FtsH periplasmic domain in novel orientation (NO), EMD-66015 (FtsH transmembrane domain), EMD-66014 (FtsH transmembrane and periplasmic domain), and in the Protein Data Bank (PDB) as PDB entry 9WUS (FtsH periplasmic domain).

## Acknowledgments

This study was funded by the Scientific and Technological Council of Türkiye (TÜBİTAK) (Project No: 118C225). G.G., and B.V.K. were funded by TÜBİTAK BİDEB 2232 International Outstanding Researchers Program (Project No: 118C225). G.G. was also funded by TÜBİTAK 2211-A National Graduate Fellowship Program. C.S. acknowledges funding by a BBSRC Responsive Mode Grant (BB/P000940/1) and a Wellcome Trust Investigator Grant (210701/Z/18/Z). The authors acknowledge support and assistance by the Wolfson Bioimaging Facility and the GW4 Facility for High-Resolution Electron Cryo-Microscopy funded by the Wellcome Trust (202904/Z/16/Z and 206181/Z/17/Z) and BBSRC (BB/R000484/1). The authors thank Mehmet Çalıseki for his help in AlphaFold2 model of the hexameric FtsH.

## Author contributions

B.V.K. and C.S. conceived and designed experiments. G.O. and G.G. purified proteins and performed *in vitro* activity assays. S.K.N.Y. performed NS-EM and cryo-EM sample preparation and preliminary image processing. U.B. collected cryo-EM data. B.V.K. and G.G. performed 3D reconstruction, and atomic model building. G.G., C.S., I.B. and B.V.K. wrote the manuscript with input from all authors.

## References

Akkulak, H., İnce, H. K., Goc, G., Lebrilla, C. B., Kabasakal, B. V., & Ozcan, S. (2024). Structural proteomics of a bacterial mega membrane protein complex: FtsH-HflK-HflC. International Journal of Biological Macromolecules, 269, 131923. 10.1016/j.ijbiomac.2024.131923

Bieniossek, C., Nie, Y., Frey, D. et al. Automated unrestricted multigene recombineering for multiprotein complex production. Nat Methods 6, 447–450 (2009a). 10.1038/nmeth.1326

Bieniossek, C., Niederhauser, B., & Baumann, U. M. (2009b). The crystal structure of apo-FtsH reveals domain movements necessary for substrate unfolding and translocation. Proceedings of the National Academy of Sciences of the United States of America, 106(51), 21579–21584. 10.1073/pnas.0910708106

Bieniossek, C., Schalch, T., Bumann, M., Meister, M., Meier, R., & Baumann, U. (2006). The molecular architecture of the metalloprotease FtsH. Proceedings of the National Academy of Sciences of the United States of America. 10.1073/pnas.0600031103

Bittner, L.-M., Arends, J., & Narberhaus, F. (2016). Mini review: ATP-dependent proteases in bacteria. Biopolymers, 105(8), 505–517. 10.1002/bip.22831

Bittner, L.-M., Arends, J., & Narberhaus, F. (2017). When, how and why? Regulated proteolysis by the essential FtsH protease in Escherichia coli. Biological Chemistry, 398(5–6), 625–635. doi:10.1515/hsz-2016-0302

Burnley, T., Palmer, C.-M., Winn, M. (2017). Recent developments in the CCP-EM software suite. Acta Crystallogr D Struct Biol. 1;73(Pt 6):469–477. doi: 10.1107/S2059798317007859.

Bruckner, R. C., Gunyuzlu, P. L., & Stein, R. L. (2003). Coupled kinetics of ATP and peptide hydrolysis by Escherichia coli FtsH protease. Biochemistry, 42(36), 10843–10852. 10.1021/bi034516h

Carvalho, V., Prabudiansyah, I., Kovacik, L., Chami, M., Kieffer, R., van der Valk, R., de Lange, N., Engel, A., & Aubin-Tam, M.-E. (2021). The cytoplasmic domain of the AAA+ protease FtsH is tilted with respect to the membrane to facilitate substrate entry. Journal of Biological Chemistry, 296. 10.1074/jbc.RA120.014739

Chen, I., Pant, S., Wu, Q., Cater, R. J., Sobti, M., Vandenberg, R. J., Stewart, A. G., Tajkhorshid, E., Font, J., & Ryan, R. M. (2021). Glutamate transporters have a chloride channel with two hydrophobic gates. Nature, 591(7849), 327–331. 10.1038/s41586-021-03240-9

Chiba, S., Akiyama, Y., & Ito, K. (2002). Membrane protein degradation by FtsH can be initiated from either end. Journal of Bacteriology, 184(17), 4775–4782. 10.1128/JB.184.17.4775-4782.2002

Chiba, S., Akiyama, Y., Mori, H., Matsuo, E., & Ito, K. (2000). Length recognition at the N-terminal tail for the initiation of FtsH-mediated proteolysis. EMBO Reports, 1(1), 47–52. 10.1093/embo-reports/kvd005

Emsley, P., Lohkamp, B., Scott, W. G., & Cowtan, K. (2010). Features and development of Coot. Acta crystallographica. Section D, Biological crystallography, 66(Pt 4), 486–501. 10.1107/S0907444910007493

Fernandez-Leiro, R., & Scheres, S. H. W. (2017). A pipeline approach to single-particle processing in RELION. Acta Crystallographica. Section D, Structural Biology, 73(Pt 6), 496–502. 10.1107/S2059798316019276

Ghanbarpour, A., Telusma, B., Powell, B. M., Zhang, J. J., Bolstad, I., Vargas, C., Keller, S., Baker, T., Sauer, R. T., & Davis, J. H. (2025). An asymmetric nautilus-like HflK/C assembly controls FtsH proteolysis of membrane proteins. The EMBO Journal: 1–13. 10.1038/s44318-025-00408-1

Ito, K., & Akiyama, Y. (2005). Cellular functions, mechanism of action, and regulation of FtsH protease. Annual Review of Microbiology, 59, 211–231. 10.1146/annurev.micro.59.030804.121316

Jumper, J., Evans, R., Pritzel, A. et al. (2021). Highly accurate protein structure prediction with AlphaFold. Nature 596, 583–589. 10.1038/s41586-021-03819-2

Kato, Y., & Sakamoto, W. (2018). FtsH protease in the thylakoid membrane: Physiological functions and the regulation of protease activity. Frontiers in Plant Science, 9. 10.3389/fpls.2018.00855

Kihara, A., Akiyama, Y., & Ito, K. (1996). A protease complex in the Escherichia coli plasma membrane: HflKC (HflA) forms a complex with FtsH (HflB), regulating its proteolytic activity against SecY. The EMBO journal, 15(22), 6122–6131.

Kihara, A., Akiyama, Y., Ito, K. (1998). Dislocation of membrane proteins in FtsH-mediated proteolysis. EMBO J. 18(11):2970–2981. doi: 10.1093/emboj/18.11.2970

Krzywda, S., Brzozowski, A.M., Verma, C., Karata, K., Ogura, T., Wilkinson, A.J. (2002). The crystal structure of the AAA domain of the ATP-dependent protease FtsH of Escherichia coli at 1.5 A resolution. Structure. Aug;10(8):1073–83. doi: 10.1016/s0969-2126(02)00806-7.

Li, X., Mooney, P., Zheng, S., Booth, C. R., Braunfeld, M. B., Gubbens, S., Agard, D. A., & Cheng, Y. (2013). Electron counting and beam-induced motion correction enable near-atomic-resolution single-particle cryo-EM. Nature Methods, 10(6), 584–590. 10.1038/nmeth.2472

Liebschner, D., Afonine, P. V., Baker, M. L., Bunkóczi, G., Chen, V. B., Croll, T. I., Hintze, B., Hung, L. W., Jain, S., McCoy, A. J., Moriarty, N. W., Oeffner, R. D., Poon, B. K., Prisant, M. G., Read, R. J., Richardson, J. S., Richardson, D. C., Sammito, M. D., Sobolev, O. V., Stockwell, D. H., … Adams, P. D. (2019). Macromolecular structure determination using X-rays, neutrons and electrons: recent developments in Phenix. Acta crystallographica. Section D, Structural biology, 75(Pt 10), 861–877. 10.1107/S2059798319011471

Liu, W., Schoonen, M., Wang, T., McSweeney, S., & Liu, Q. (2022). Cryo-EM structure of transmembrane AAA+ protease FtsH in the ADP state. Communications Biology, 5(1). 10.1038/s42003-022-03213-2

Ma, C., Wang, C., Luo, D., Yan, L., Yang, W., Li, N., & Gao, N. (2022a). Structural insights into the membrane microdomain organization by SPFH family proteins. Cell Research, 32(2), 176–189. 10.1038/s41422-021-00598-3

Meng, E. C., Goddard, T. D., Pettersen, E. F., Couch, G. S., Pearson, Z. J., Morris, J. H., & Ferrin, T. E. (2023). UCSF ChimeraX: Tools for structure building and analysis. Protein Science□: A Publication of the Protein Society, 32(11), e4792. 10.1002/pro.4792

Mindell, J. A., & Grigorieff, N. (2003). Accurate determination of local defocus and specimen tilt in electron microscopy. Journal of Structural Biology, 142(3), 334–347. 10.1016/s1047-8477(03)00069-8

Mirdita, M., Schütze, K., Moriwaki, Y., Heo, L., Ovchinnikov, S., & Steinegger, M. (2022). ColabFold: Making protein folding accessible to all. Nature Methods, 19(6), 679–682. 10.1038/s41592-022-01488-1

Murata, K., & Wolf, M. (2018). Cryo-electron microscopy for structural analysis of dynamic biological macromolecules. Biochimica et Biophysica Acta - General Subjects, 1862(2), 324–334. 10.1016/j.bbagen.2017.07.020

Murshudov, G. N., Skubák, P., Lebedev, A. A., Pannu, N. S., Steiner, R. A., Nicholls, R. A., Winn, M. D., Long, F., & Vagin, A. A. (2011). It REFMAC5 for the refinement of macromolecular crystal structures. Acta Crystallographica Section D, 67(4), 355–367. 10.1107/S0907444911001314

Pettersen, E. F., Goddard, T. D., Huang, C. C., Meng, E. C., Couch, G. S., Croll, T. I., Morris, J. H., & Ferrin, T. E. (2021). UCSF ChimeraX: Structure visualization for researchers, educators, and developers. Protein science : a publication of the Protein Society, 30(1), 70–82. 10.1002/pro.3943

Puchades, C., Rampello, A. J., Shin, M., Giuliano, C. J., Wiseman, R. L., Glynn, S. E., & Lander, G. C. (2017). Structure of the mitochondrial inner membrane AAA+ protease YME1 gives insight into substrate processing. Science (New York, N.Y.), 358(6363). 10.1126/science.aao0464

Qiao, Z., Yokoyama, T., Yan, X.-F., Beh, I. T., Shi, J., Basak, S., Akiyama, Y., & Gao, Y.-G. (2022). Cryo-EM structure of the entire FtsH-HflKC AAA protease complex. Cell Reports, 39(9), 110890. 10.1016/j.celrep.2022.110890

Rosenthal, P. B., & Henderson, R. (2003). Optimal determination of particle orientation, absolute hand, and contrast loss in single-particle electron cryomicroscopy. Journal of Molecular Biology, 333(4), 721–745. 10.1016/j.jmb.2003.07.013

Sauer, R. T., & Baker, T. A. (2011). AAA+ proteases: ATP-fueled machines of protein destruction. Annual Review of Biochemistry, 80, 587–612. 10.1146/annurev-biochem-060408-172623

Sanchez-Garcia, R., Gomez-Blanco, J., Cuervo, A. et al. DeepEMhancer: a deep learning solution for cryo-EM volume post-processing. Commun Biol 4, 874 (2021). 10.1038/s42003-021-02399-1

Scharfenberg, F., Serek-Heuberger, J., Coles, M., Hartmann, M. D., Habeck, M., Martin, J., Lupas, A. N., & Alva, V. (2015). Structure and Evolution of N-domains in AAA Metalloproteases. Journal of Molecular Biology, 427(4), 910–923. 10.1016/j.jmb.2014.12.024

Suno, R., Niwa, H., Tsuchiya, D., Zhang, X., Yoshida, M., & Morikawa, K. (2006). Structure of the whole cytosolic region of ATP-dependent protease FtsH. Molecular Cell, 22(5), 575–585. 10.1016/j.molcel.2006.04.020

Vostrukhina, M., Popov, A., Brunstein, E., Lanz, M., Baumgartner, R., Bieniossek, C., Schacherl, M., & Baumann, U. (2015). The structure of Aquifex aeolicus FtsH in the ADP-bound state reveals a C 2 -symmetric hexamer. Acta Crystallographica Section D Biological Crystallography, 71. 10.1107/S1399004715005945

Wang, S., Wang, K., Song, K., Lai, Z. W., Li, P., Li, D., Sun, Y., Mei, Y., Xu, C., & Liao, M. (2024). Structures of the Mycobacterium tuberculosis efflux pump EfpA reveal the mechanisms of transport and inhibition. Nature Communications, 15(1), 7710. 10.1038/s41467-024-51948-9

Yamada-Inagawa, T., Okuno, T., Karata, K., Yamanaka, K., & Ogura, T. (2003). Conserved Pore Residues in the AAA Protease FtsH Are Important for Proteolysis and its Coupling to ATP Hydrolysis. Journal of Biological Chemistry, 278(50), 50182–50187. 10.1074/jbc.M308327200

Yedidi, R. S., Wendler, P., & Enenkel, C. (2017). AAA-ATPases in protein degradation. Frontiers in Molecular Biosciences, 4(JUN). 10.3389/fmolb.2017.00042

Yi, L., Liu, B., Nixon, P. J., Yu, J., & Chen, F. (2022). Recent Advances in Understanding the Structural and Functional Evolution of FtsH Proteases. Frontiers in Plant Science, 13. 10.3389/fpls.2022.837528

Zivanov, J., Nakane, T., & Scheres, S. H. W. (2020). Estimation of high-order aberrations and anisotropic magnification from cryo-EM data sets in RELION-3.1. IUCrJ, 7(Pt 2), 253–267. 10.1107/S2052252520000081

