## Supplementary Information for "Cryo-EM structure of the FtsH periplasmic domain reveals functional dynamics"

[‡] Present address: University Medical Center Hamburg-Eppendorf, Hamburg, Germany

* To whom correspondence may be addressed

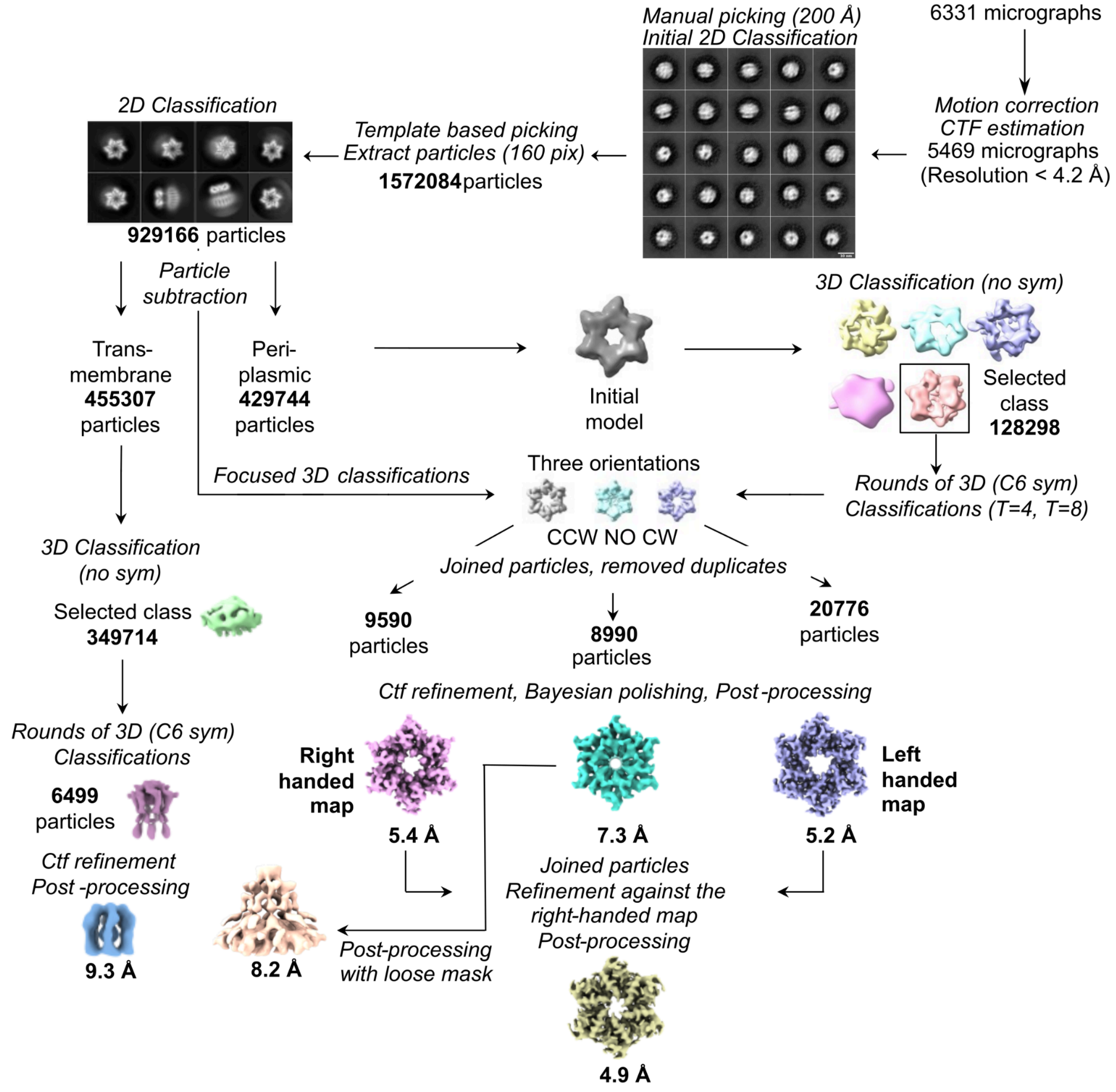


**Figure S1. Cryo-EM workflow.** FtsH periplasmic domain (PD) and transmembrane domain (TM) sub-regions were reconstructed separately by using subtracted particles extracted with masks of corresponding regions. The FtsH-PD region was reconstructed by focused 3D classification. Three different orientations were obtained for FtsH-PD; two of them (right-hand and left-hand) belong to a known FtsH-PD conformation, the third map exhibits a novel conformation (NO: novel orientation). The particles corresponding to left-handed (CW) and right-handed (CCW) maps were combined and refined to obtain the right-handed map. PD and TM regions were obtained in one map by applying a loose mask to the NO map of FtsH-PD.

**Table S1. Cryo-EM Data Collection**

| | |
|---|---|
| **Voltage (kV)** | 200 |
| **Nominal Magnification** | 130,000 |
| **Pixel Size [Å]** | 1.05 (0.525) |

| | |
|---|---|
| **Total Dose [e⁻/ Å²]** | 61.3 |
| **Number of Fractions** | 48 |
| **Total Dose per Fraction [e⁻/ Å²]** | 1.28 |
| **Defocus Range [μm]** | -1 to -2 |
| **No. of micrographs** | 6331 |
| **Initial particle images** | 929,166 |
| **Final particle images** | 30,366 |
| **Symmetry imposed** | C6 |

**Table S2. FtsH-PD Refinement**

| | |
|---|---|
| **Initial mode used (PDB)** | 7WI3 |
| **PDB code** | 9WUS |
| **EMDB code** | EMD-66269 |
| **Composition** | |
| **Chains** | 6 |
| **Atoms** | 3168 |
| **Residues** | 390 |
| **Water** | 0 |
| **Ligands** | 0 |
| **Bonds (RMSD)** | |
| **Length (Å)** | 0.002 |
| **Angles (°)** | 0.567 |
| **B-factor (Å²)** | 125.9 |
| **MolProbity score** | 2.09 |
| **Clash score** | 12.5 |
| **Ramachandran plot (%)** | |
| **Outliers** | 0 |
| **Allowed** | 7.9 |
| **Favored** | 93.65 |
| **Rotamer outliers (%)** | 0 |
| **CaBLAM outliers (%)** | 4.9 |
| **Resolution FSC (0.143)(Å)** | 4.24 |
| **CC (mask)** | 0.65 |

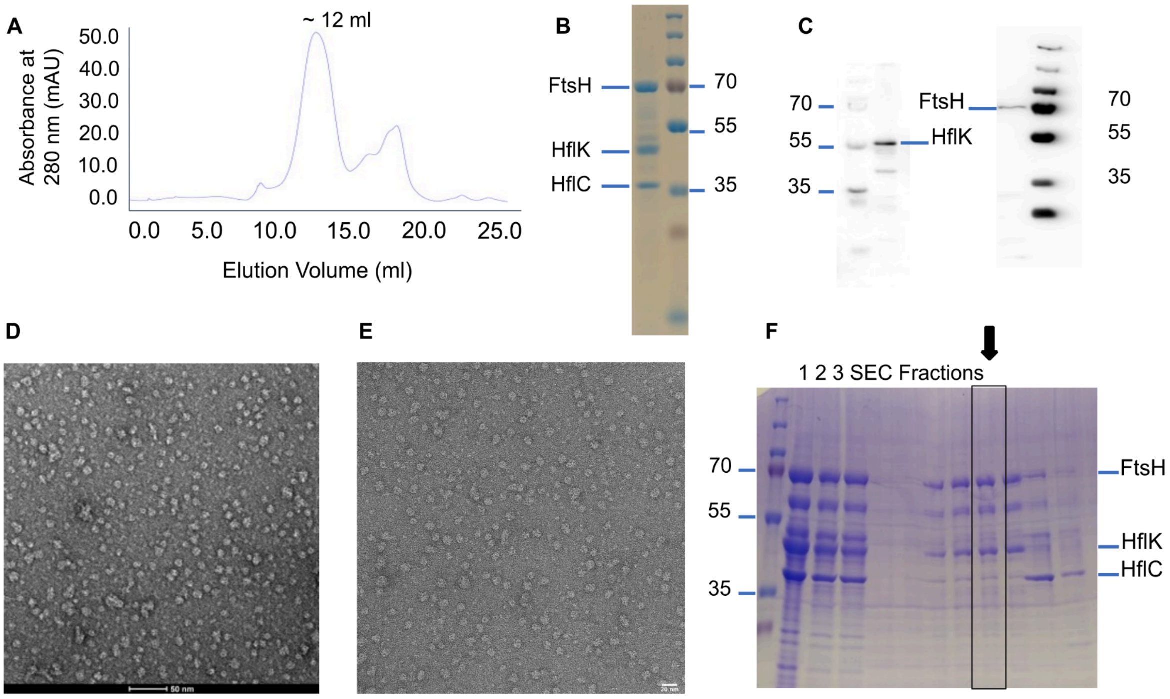


**Figure S2. Protein purification and sample preparation for EM. A.** Size-exclusion chromatography (SEC) of purified FtsH-HflKC complex. **B.** SDS-PAGE analysis of the peak fraction. **C.** Western Blot analyses of streptactin affinity-purified FtsH-HflKC. Anti-His and anti-Strep tag antibodies were used. **D.** NS-EM micrograph (Scale bar is 50 nm). **E.** Cryo-EM micrograph (Scale bar is 20 nm). **F.** SDS-PAGE analysis of Amphipol-reconstituted FtsH-HflKC. 1. After Streptactin affinity purification, 2. After complex reconstitution in Amphipol, 3. After detergent removal with Biobeads, SEC Fractions: Peak fractions during SEC. The highlighted fraction (box) was used for NS-EM and cryo-EM analysis.

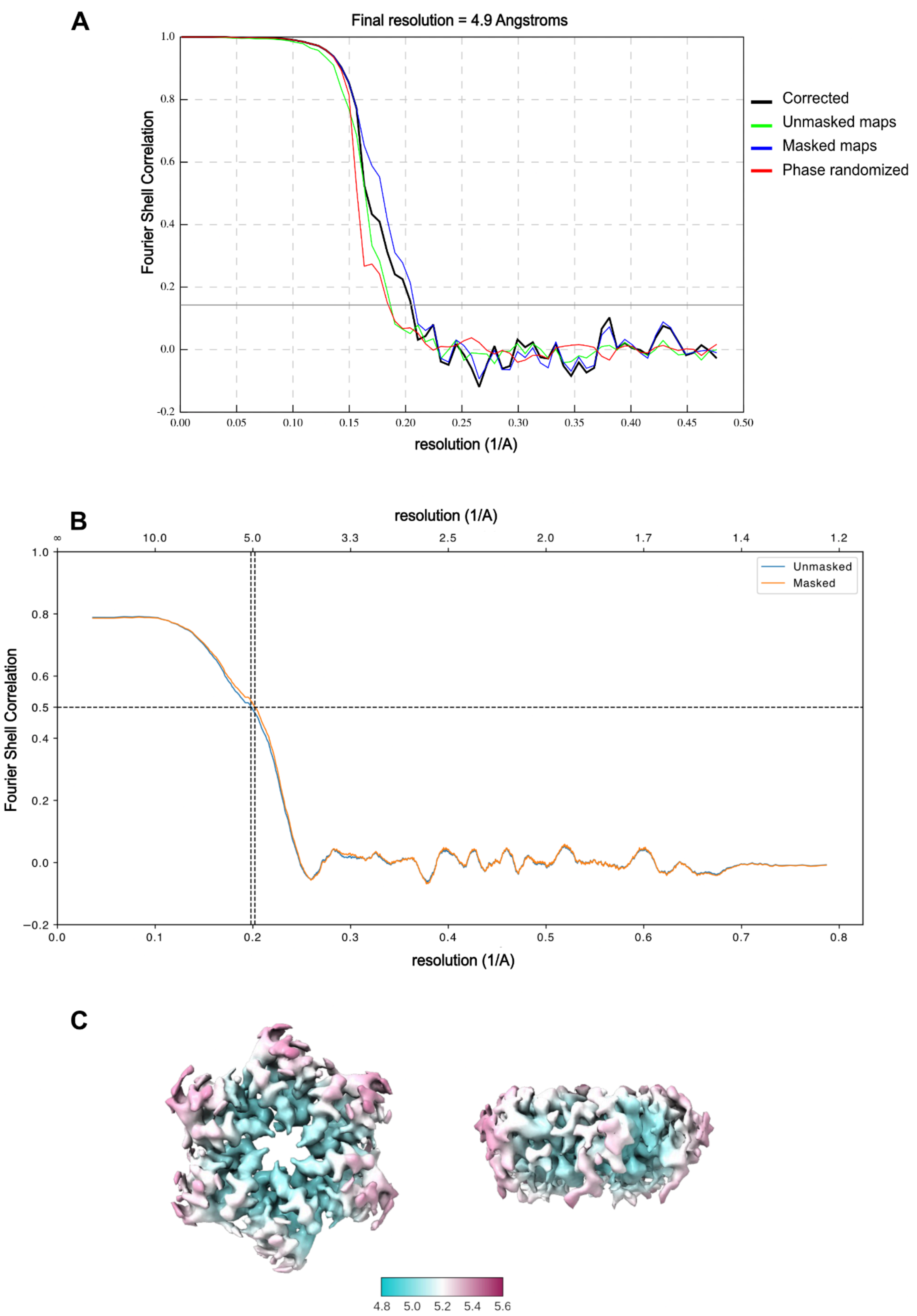


**Figure S3. The resolution of the FtsH-PD map according to FSC. A.** Gold standard Fourier Shell Correlation (FSC) curves of the FtsH-PD maps. The 4.9 Å resolution was determined using the FSC=0.143 criterion (shown in gray line). **B.** The model map FtsH-PD FSC curve calculated between the model and the map. The resolution indicates 5.0 Å at FSC=0.5. **C.** Local resolution map of FtsH-PD, varying between 4.8-5.6 Å.

A

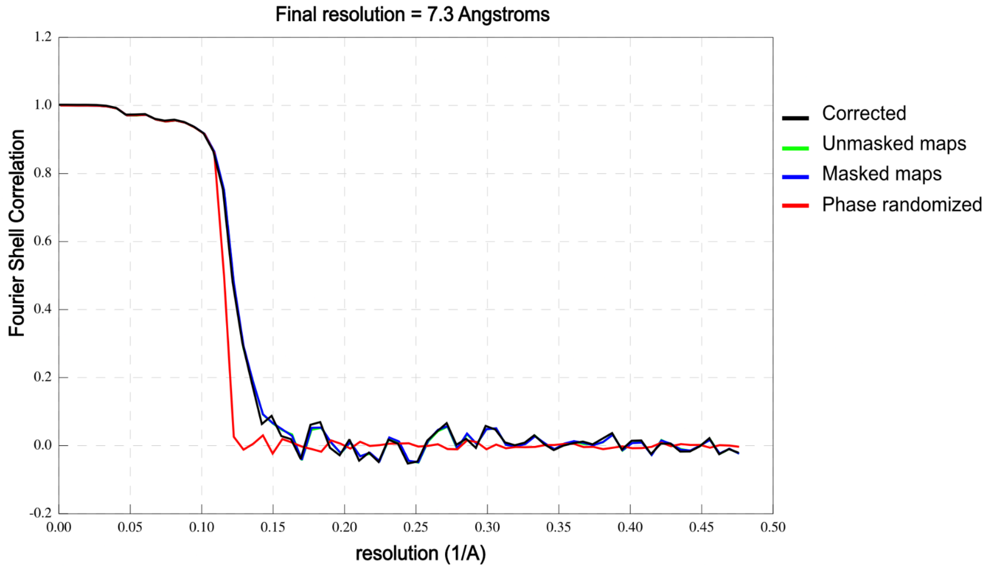


B

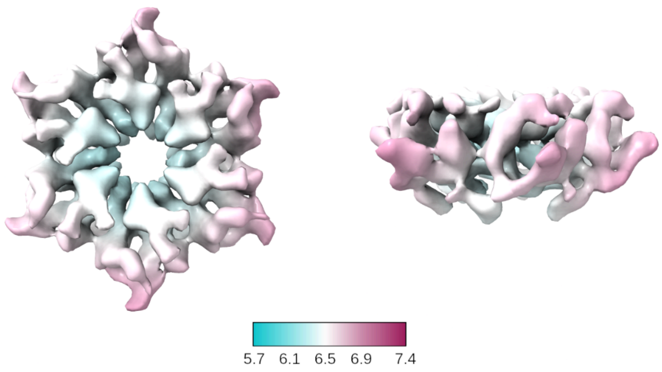


**Figure S4. The resolution of the FtsH-PD-NO map according to FSC. A.** Gold standard Fourier Shell Correlation (FSC) curves of the FtsH-PD maps. The 7.3 Å resolution was determined using the FSC=0.143 criterion **B.** Local resolution map of FtsH-PD-NO, varying between 5.7-7.4 Å.

**A**

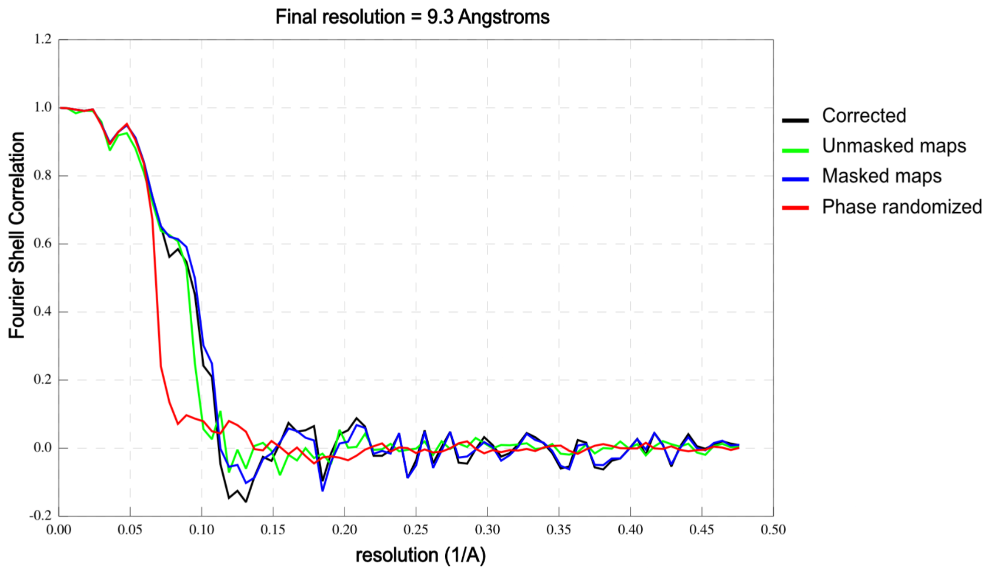


**B**

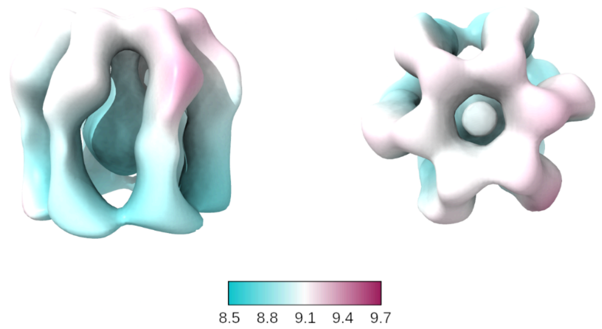


**Figure S5. The resolution of the transmembrane domain from the FtsH-PD map according to FSC. A.** Gold standard Fourier Shell Correlation (FSC) curves of the FtsH-PD maps. The 9.3 Å resolution was determined using the FSC=0.143 criterion **B.** Local resolution map of FtsH-PD-NO, varying between 8.5-9.7 Å.

**A**

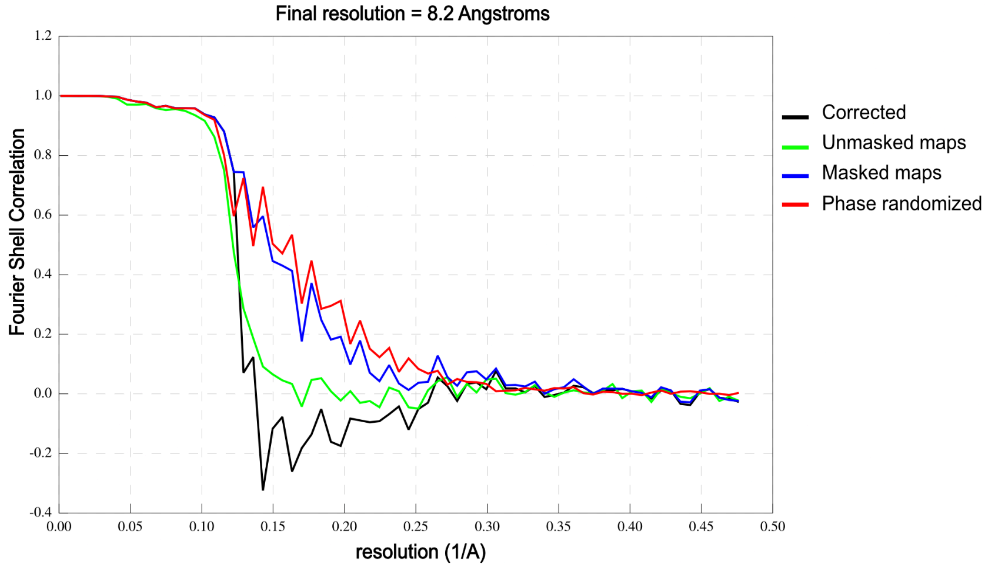


**B**

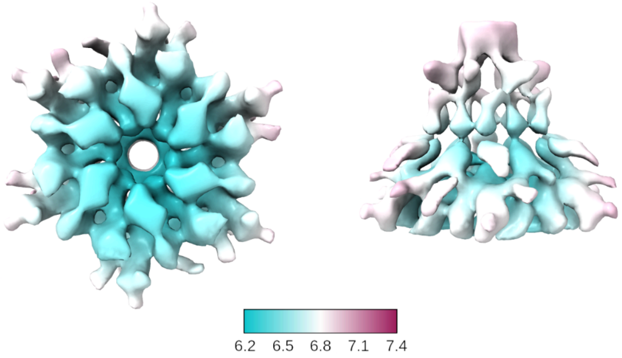


**Figure S6. The resolution of the transmembrane and periplasmic domains from FtsH-PD-NO map according to FSC. A.** Gold standard Fourier Shell Correlation (FSC) curves of the FtsH-PD maps. The 8.2 Å resolution was determined using the FSC=0.143 criterion **B.** Local resolution map of FtsH-PD-NO, varying between 6.2-7.4 Å.

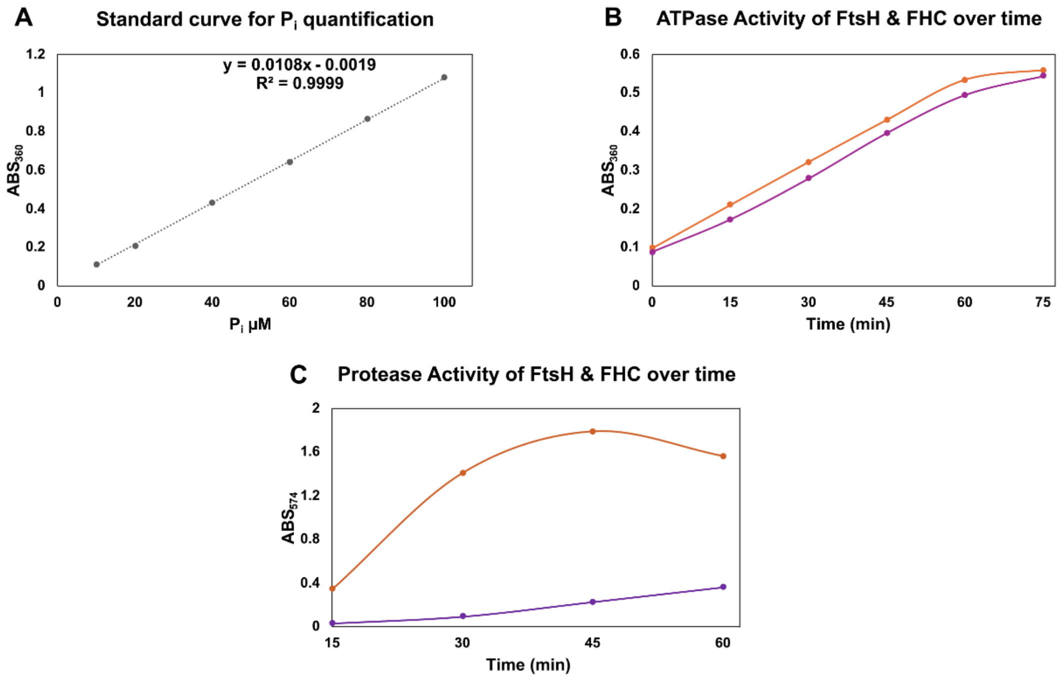


**Figure S7. ATPase and protease activities of FtsH-HflKC. A.** ATP hydrolysis was measured by quantifying inorganic phosphate ($P_i$) release over time, **B.** FtsH (orange) and FtsH-HflKC (FHC, purple) were incubated with 5 mM ATP, and $P_i$ release was monitored, then subtracted from standard curve of $P_i$ at the indicated time points. **C.** Protease activity of FtsH (orange) and FHC (purple) proteins using casein (0.4%) as substrate.

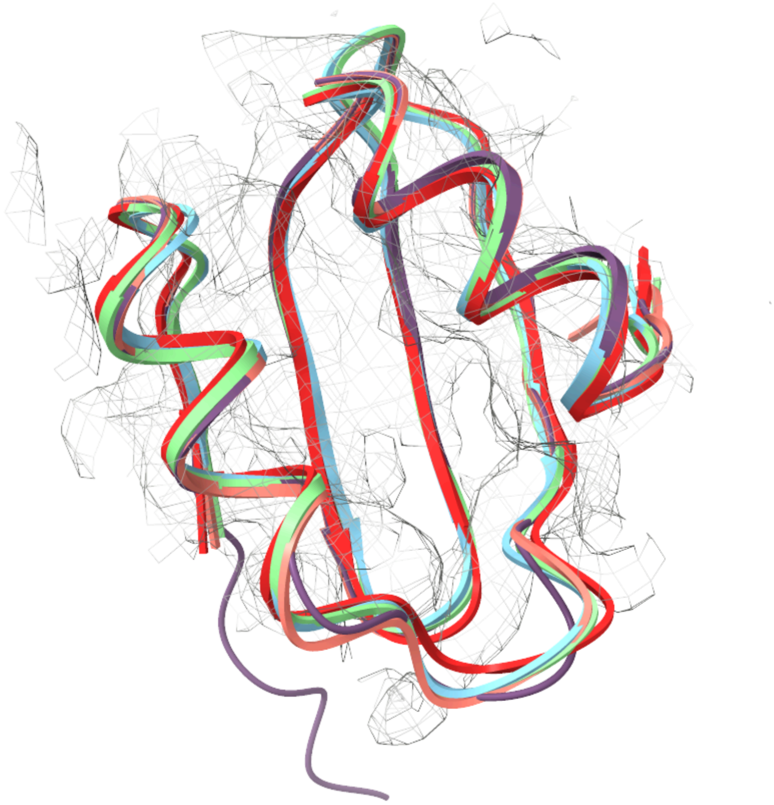

**Figure S8. Superposition of *E. coli* FtsH periplasmic domain structures.** NMR (pdb 2muY, red), X-ray (pdb 4V0B, turquoise), and cryo-EM structures (pdb 7wi3 and 7vhp, salmon and green, respectively) of FtsH periplasmic domains are aligned with the FtsH-PD structure (purple). FtsH-PD EM density is shown in grey mesh. All structures reveal the conserved α+β fold.
